# Edit Distance Embedding with Genomic Large Language Model

**DOI:** 10.1101/2025.09.25.678635

**Authors:** Xiang Li, Ke Chen, Yijia Zhang, Mingfu Shao

## Abstract

Edit distance is a fundamental metric in genomic sequence analysis, yet it is computationally expensive to calculate. A practical approach for large-scale sequence analysis involves mapping sequences into a normed space and approximating the edit distance using the more efficiently computed distance in that space. This process, known as edit distance embedding, has been extensively studied both theoretically and in practice. Recently, embedding methods based on machine learning have gained popularity, where the mapping is represented as a neural network whose parameters are learned from data. However, the accuracy of these methods remains unsatisfactory, leaving much room for improvement. Recent advancements in genomic language models have shown remarkable performance in various sequence analysis applications. We investigate if improved embeddings can be achieved using DNA language models. We introduce LLMED, a model designed to produce sequence embeddings approximating the edit distance. LLMED is trained via contrastive learning based on a pretrained genomic large language model. Through extensive experimental comparisons, we show that LLMED surpasses leading machine learning and rule-based embedding methods in approximating the edit distance; LLMED also achieved significantly improved accuracy in a critical application, similar sequence search.

## 1 Background

Edit distance (Levenshtein distance) is an elementary measure for comparing genomic sequences. Classical methods compute the exact edit distance and the optimal alignment, but their quadratic time complexity makes it unsuitable for large-scale applications. Approximation algorithms for the edit distance has been an active area of theoretical research, resulting in superlinear (but sub-quadratic) algorithms with constant approximation ratios and linear-time algorithms with polynomial approximation ratios [2, 3]. These algorithms remain largely of theoretical interests primarily due to their large approximation factors. Practical methods and implementations such as Wavefront algorithm [20], A*PA2 [12], and edlib [36] have improved the efficiency through algorithmic innovations and hardware optimizations. However they often rely on the assumption that the resulting edit distance is small. Seeding strategies such as k-mers and strobemers [26, 27], combined with sketching techniques including minimizers [28, 25] and syncmers [11], have been developed to estimate the Average Nucleotide Identity (ANI) and mutation rates by comparing the lightweight sketches [29, 24]; nevertheless they are not intended to approximate the edit distance.

Recently, embedding-based methods have gained prominence as an efficient approach for sequence similarity estimation. After embedding sequences into a metric space, typically a normed space such as Hamming distance or Euclidean distance, the edit distance can be estimated by computing a distance in the embedded space which is often simpler and faster. This approach enables broad applications such as nearest sequence search [8, 6], alignment-free similarity estimation [14], and phylogeny reconstruction [19]. Embedding has been studied in theory, where one seeks an embedding with low distortion [23]. It has been proved that, the edit distance cannot be embedded into *l*_1_ norm with a distortion better than 3*/*2 [1]. On the positive side, CGK [5] embeds the edit distance into the hamming space using a randomized injective approach and is proved to admit a distortion of *O*(*k*^2^) where *k* denotes the edit distance. Heuristic embedding methods have been developed, which can be classified as rule-based methods and machine learning methods [6]. Some rule-based approaches are using substrings, includes FFP [30], which utilizes the frequency profile of fixed-length substrings, and Mash [14], which applies MinHash technique [4] on substrings. Smooth-q [31] was developed for detecting overlaps among error-prone long reads based on CGK embeddings. Tensor Sketching [17, 16], which generates fixed-length embeddings by partitioning and counting subsequences, has been developed for estimating edit distance, inferring phylogenies, and aligning sequences to graphs.

Learning-based methods include a two-layer neural network with gated recurrent units (GRU) structure [34], trained using a three-stage training process with distinct loss functions at each stage. CNNED [8] demonstrated that the convolutional neural network (CNN) structure is more effective for the edit distance embedding than recurrent neural network (RNN) models, achieving superior performance in nearest neighbor search for biological sequences. NeuroSEED [7] employed global and local transformers to encode biological sequences, outperforming other models in tasks such as edit distance approximation, hierarchical clustering, and multiple sequence alignment. Bio-kNN [6] proposed a sequence k-nearest neighbor (KNN) search framework that combines a multihead CNN model with a specially designed triplet data selection strategy achieving state-of-the-art performance in the KNN search task.

Large language models (LLMs), also known as foundation models, initially emerged in natural language processing [32, 10], have recently gained popularity in genomic sequence analysis. These models process long input sequences by tokenizing them and calculating embeddings using architectures such as Transformers. LLMs are typically pretrained on large datasets and fine-tuned for various downstream tasks. DNABERT [15] adapted the Bidirectional Encoder Representations from Transformers (BERT) framework to train on the human genome, enabling applications such as identifying transcription factor binding sites, splicing sites, and genetic variants. Nucleotide Transformer [9] improved the scalability of the genomic foundation models by training on genomes from 850 species and replacing overlapping k-mer tokenization with a non-overlapping approach, largely reducing tokenized sequence lengths. DNABERT2 [35] further enhanced tokenization with Byte Pair Encoding technique, outperforming other models on the GUE benchmark. Beyond Transformer architectures, alternative LLM designs have also been explored. Evo [21] and HyenaDNA [22] employ the (stripped) Hyena architecture to efficiently handle genome-scale input sequences. Notably, Evo enables both prediction and generation of DNA sequences. Mamba [13], which employs structured state space models (SSMs) that have been developed to address Transformers’ computational inefficiency on long sequences, has been applied to species classification tasks with genome fragments as inputs.

In this paper, we explore if genomic language models can be used to accurately approximate edit distance, a problem that has not been previously explored. We present LLMED, a genomic sequence embedding model that explicitly designed to generate high-quality embeddings that approximate edit distances. Our model outperforms other sequence embedding models in edit distance estimation and achieves state-of-the-art performance in top-*K* similar sequence searches. By addressing limitations in current approaches, our work establishes a new benchmark for sequence embedding in genomic analysis.

## 2 Results and Discussion

We compare our genomic sequence embedding model, LLMED, against established embedding baselines. Each embedding model maps an input genomic sequence to a fixed-dimensional vector. We first quantify the correlation between each model’s embedding distance and the true edit distance (Section 2.1). We then evaluate if the edit distance can be approximated from the embedding distance and assess their accuracies (Section 2.2). Finally, we examine practical utility in a real-world setting via *K*-nearest-neighbor sequence search (Section 2.3).

### 2.1 Correlation with Edit Distance

We measure the correlation between the edit distance of genomic sequences and the distance induced by each model’s embeddings. To this end, we construct a simulated evaluation dataset as follows. For each length *L* ∈ {100, 200, 500, 1000}, we generate 5000 sequence pairs over {*A, C, G, T*}. In each pair, the first sequence is a random string of length *L*; the second is obtained by introducing random substitutions, insertions, and deletions with equal probabilities. The edit distance of each simulated pair is calculated using dynamic programming as the ground-truth. We ensure that the true edit distances of the 5000 pairs are uniformly distributed in {1, 2, …, ⌊*L/*2⌋}.

We compare LLMED with several established sequence-embedding approaches. LLMED is evaluated using three different loss function variants: MAE loss, triplet loss and combined loss. All variants are initialized on the DNABERT2 pretrained model and fine-tuned on an independently generated dataset; training details are provided in Section 4. Throughout, we quantify output-space similarity using cosine distance (1− cosine similarity) between embeddings. For the baselines, we evaluate Tensor Sketch, a rule-based method that has demonstrated strong correlation with edit distance among rule-based methods [17]. We apply the Tensor Slide Sketch variant with embedding dimension of 64, window size set to 10% of the sequence length *L*, and stride set to 1% of the sequence length *L*, based on the parameter settings described in the original paper. The distance measure for Tensor Sketch is squared Euclidean norm as defined in the method’s original implementation. We also benchmark a recent machine-learning embedding method: CNNED [8]. For CNNED, we follow its training pipeline outlined in the original paper. Specifically, we train the model on 1000 randomly chosen sequences for each dataset with 50 epochs. For datasets with *L* ranging from 100 to 1000, the number of CNN layers is adjusted from 5 to 8 accordingly to make sure the model fits the sequence length. Additionally, we utilize Euclidean distance as distance measure as specified in its implementation. While Bio-KNN has been shown to achieve strong performance in its experimental results [6], it provides models only for protein sequences and is therefore excluded from our experiments. Last, we evaluate the genomic foundation model DNABERT2 [35] that is used in LLMED. For DNABERT2, the sequence embeddings are computed by applying average pooling across all token positions, excluding special tokens such as start and end markers. Consistent with our model, the cosine distance between output embeddings is used as the distance measure. We utilize the pretrained checkpoint DNABERT-2-117M without any fine-tuning.

To first provide an intuitive view of correlation, in Fig. 1 we plot the edit distance versus the embedding distance for all 5000 pairs for the case of *L* = 1000. We can observe that LLMED exhibits a clearer, tighter monotonic trend than other baselines.

**Figure 1:**
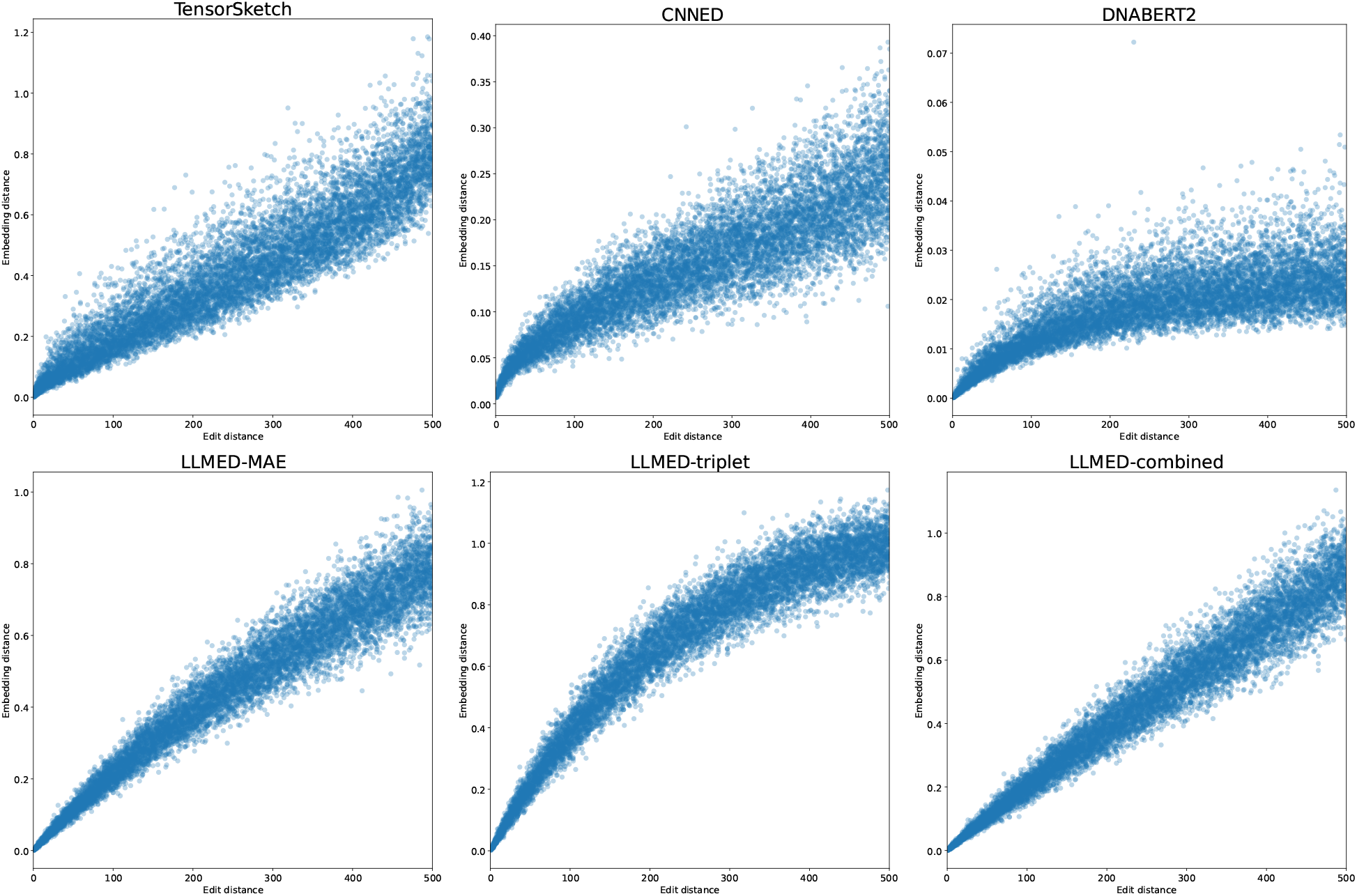
Edit distance (x-axis) versus the embedding distance (y-axis).

We then report the Spearman’s rank correlation and Pearson correlation between the embedding distance and the ground-truth edit distance. Fig. 2 illustrates the results with varying sequence lengths. We can observe that LLMED consistently outperforms other embedding models, achieving the best correlation for different *L* on both measures. In particular, at *L* = 1000, each LLMED variant exceeds 95% Spearman correlation. Pearson correlations mirror these trends, indicating that the relationship is also approximately linear. Tensor Sketch and CNNED are competitive yet remain behind LLMED, while DNABERT2 shows much poorer performance, demonstrating the importance of our training procedure in improving the model’s ability to approximate edit distance.

**Figure 2:**
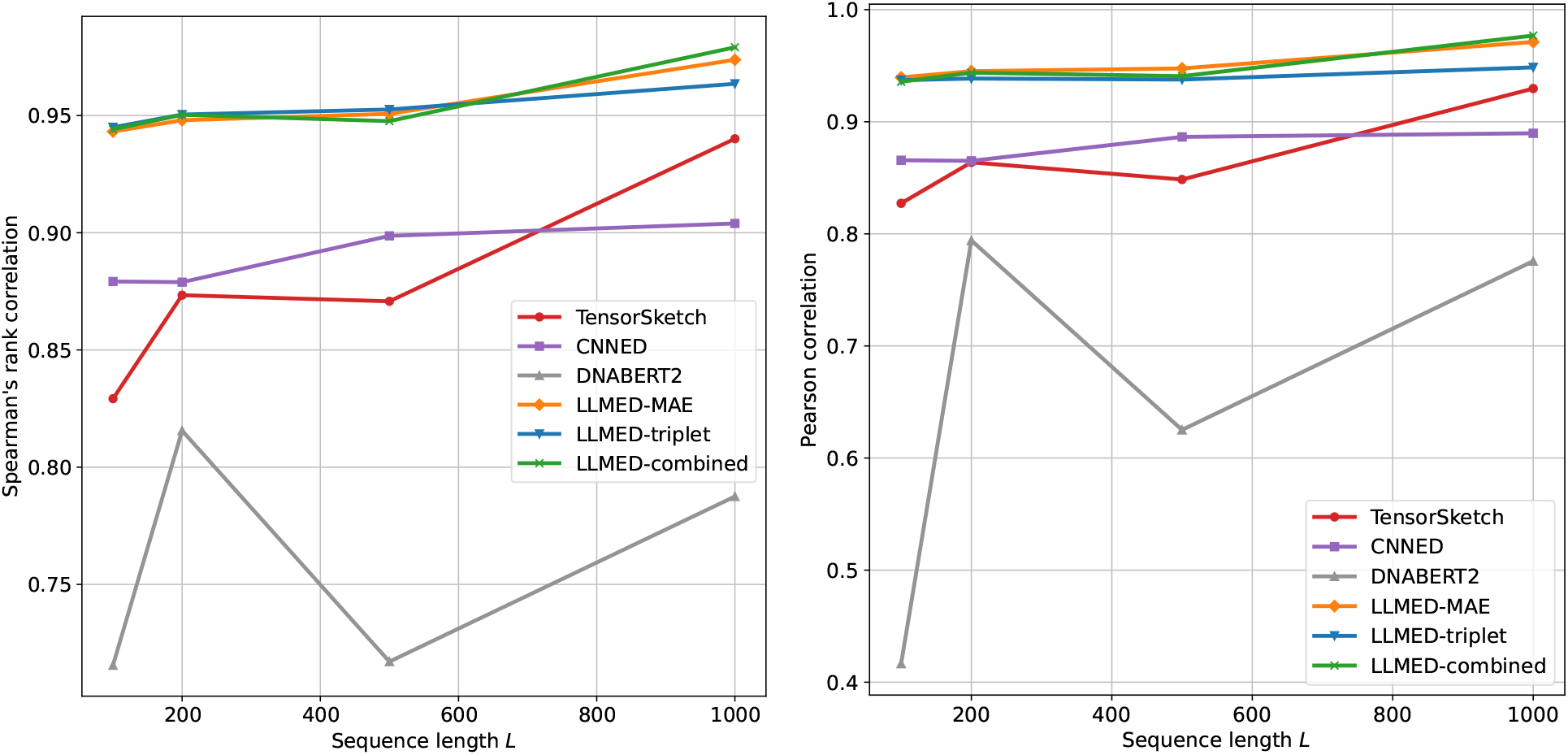
Left: Spearman’s rank correlation between the true edit distance and the embedding distance for datasets with different sequence length *L*. Right: Pearson correlation between the true edit distance and the embedding distance for datasets with different sequence length *L*.

### 2.2 Approximating the Edit Distance

Next, we evaluate how well the edit distance can be approximated from the embeddings. We examine two strategies for reconstructing the edit distance from a method’s embedding distance. (1) for each method, we fit a linear function that maps from the embedding distance to the edit distance, trained on an independently simulated calibration set; (2) for our methods (the three variants of LLMED), we also use a fixed closed-form approximation:

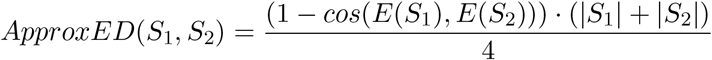

where *S*_1_ and *S*_2_ are the two input genomic sequences, cos(·, ·) denotes cosine similarity and *E*(·) is the embedding generated by our model. This approximation is derived from the loss function used to fine-tune our models; see Section 4. Note that this method does not require additional training.

We plot the true edit distance against the approximated edit distance in Fig. 3 for L = 1000 on the same set of simulated pairs used in Section 2.1. Perfect estimates would lie on the diagonal *y* = *x*. The points in LLMED figures, especially using the combined loss, fits this line better than other methods.

**Figure 3:**
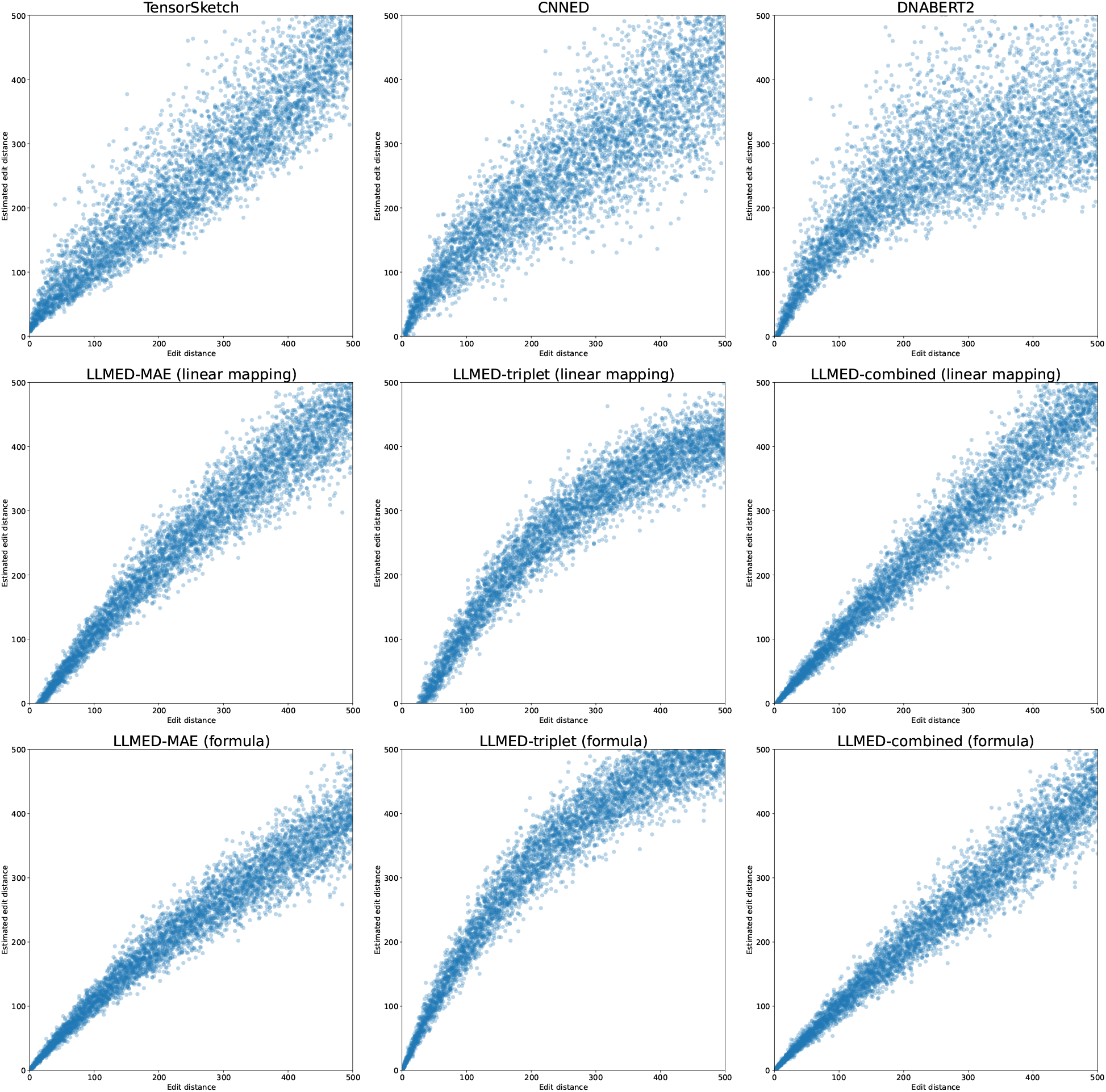
Edit distance versus estimated edit distance. For Tensor Sketch, CNNED, DNABERT2, the embedding distance is transformed to an edit distance using fitted linear mapping, whereas LLMED variants employ either a fitted linear mapping or a fixed formula.

To quantitatively compare different methods for different *L*, we report two measures. Given an evaluation set with *N* pairs of sequences, where we denote by *G*_*i*_ the true edit distance and by *A*_*i*_ the approximated edit distance by a model, 1 ≤ *i* ≤ *N*, we compute the mean squared error (MSE) and mean absolute percentage error (MAPE):

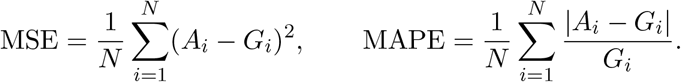

The results on MSE is shown in Fig. 4. Across different sequence lengths *L*, our methods consistently achieve the lowest error. With a fitted linear mapping (left panel), all three LLMED variants outperform Tensor Sketch, CNNED, and DNABERT2 for every *L*, where the combined-loss variant delivers the best overall accuracy. Using the fixed approximation formula (right panel) leads to a slight increase in error for the LLMED models, but LLMED-combined still attains the minimum MSE. Tensor Sketch and CNNED also perform relatively well since their designs are tailored to this task.

**Figure 4:**
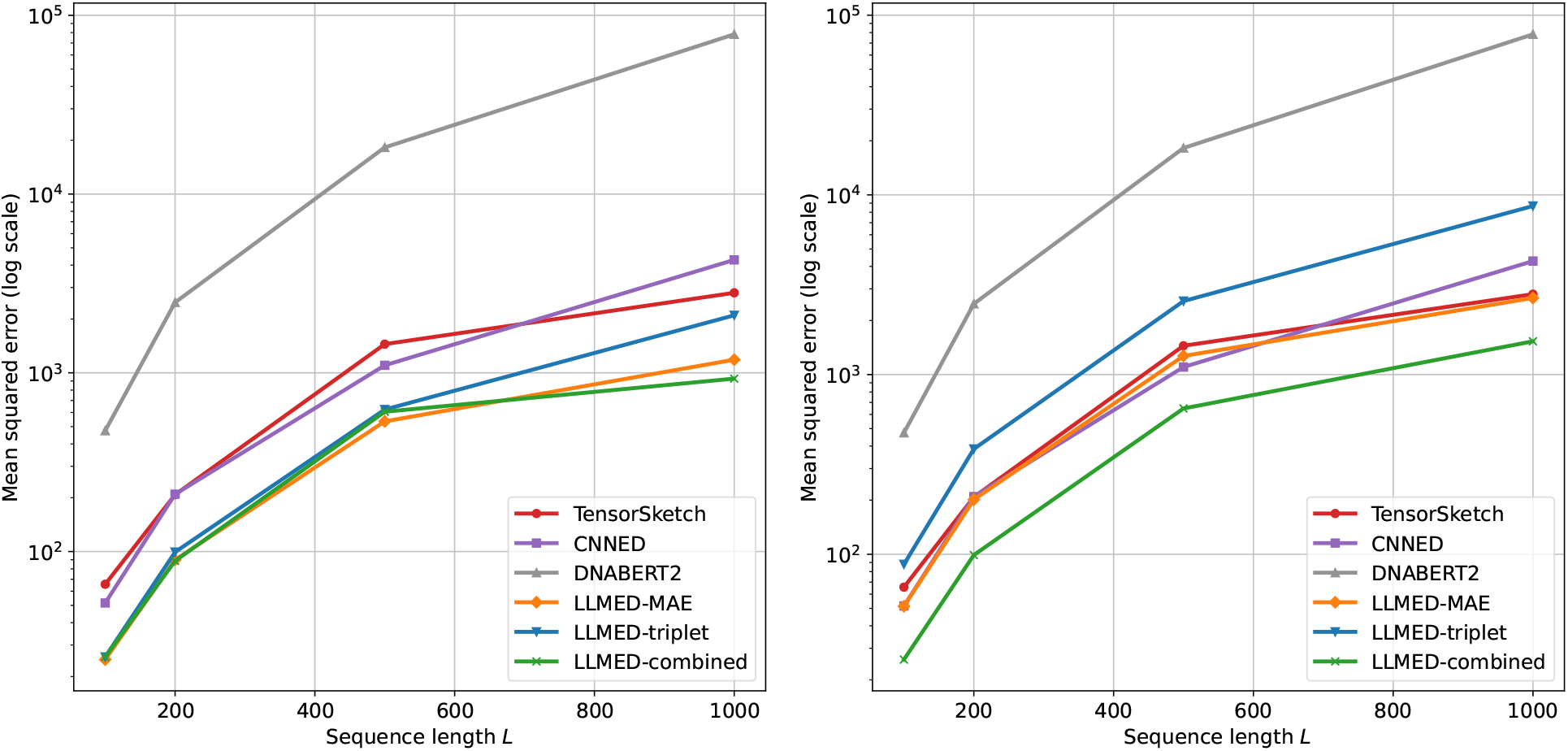
Mean squared error (MSE) versus sequence length *L* ∈ {100, 200, 500, 1000}. Left: our methods with a fitted linear mapping from the cosine similarity to edit distance. Right: our methods with the fixed approximation formula. In both panels, the baselines (Tensor Sketch, CNNED, DNABERT2) are mapped to edit distance using a fitted linear mapping.

The results reporting MAPE are given in Fig. 5. LLMED-MAE and LLMED-combined out-perform the baselines across the full range of *L*; moreover, these two models improve further when using the fixed approximation formula. Collectively, these results indicate that edit distance can be estimated accurately from our embeddings.

**Figure 5:**
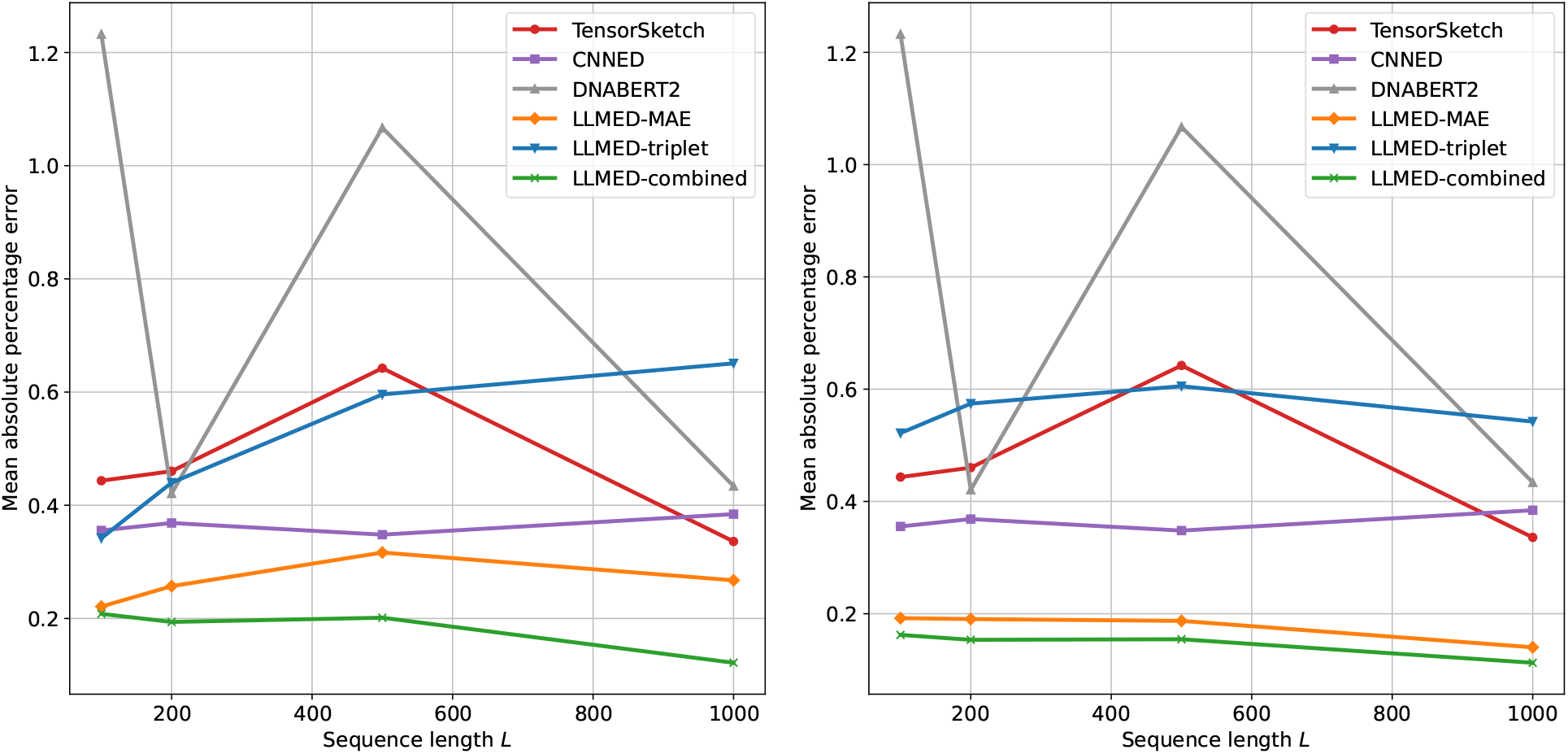
Mean absolute percentage error (MAPE) versus sequence length *L* ∈ {100, 200, 500, 1000}. Left: our methods with a fitted linear mapping from the cosine similarity to edit distance. Right: our methods with the fixed approximation formula. In both panels, the baselines (Tensor Sketch, CNNED, DNABERT2) are mapped to edit distance using a fitted linear mapping.

### 2.3 Nearest Neighbors Search

We evaluate different methods by using their embeddings on a critical application in sequence analysis, *K*-nearest neighbor search. We follow the top-*K* similar sequences search experimental setup used in CNNED [8] and Bio-KNN [6] studies. All methods are assessed on one synthetic and one real dataset. To generate the synthetic dataset, we first randomly simulate 5000 genomic sequences of length 1000. Then for each sequence, 10 similar sequences are simulated by applying random mutations with randomly selected mutation rates below 30%. Therefore, we get a synthetic dataset with 55000 sequences. The real dataset, Gen50ks [33], contains 50000 sequence fragments from 50 individual samples of human genome chromosome 20, with an average length of 5000. Both datasets are split into two parts: a query dataset with 1000 sequences, and a base dataset containing the remaining sequences. For each sequence in the query dataset, the goal is to identify the top-*K* sequences in the base dataset with the smallest edit distances. The ground-truth is obtained by computing exact edit distances using a dynamic programming algorithm between each sequence in query dataset and each sequence in base dataset.

We compare our methods with the same baselines evaluated in the previous experiments, keeping all hyperparameters and configurations identical across experiments. A sampled subset of the original dataset is used exclusively to train CNNED (8 layers for the synthetic dataset and 10 layers for Gen50ks). All LLMED variants are fine-tuned on an independently simulated dataset with sequence length *L* = 1000; no sequences from the synthetic or real benchmark datasets are used during training or fine-tuning. Given the embeddings produced by each method, we rank the sequences from base dataset by their embedding distances to a query and return the top-*K* nearest neighbors. As in previous experiments, CNNED uses Euclidean distance, Tensor Sketch uses squared Euclidean distance, and all other methods use cosine similarity.

Fig. 6 presents the results for values of *K* ranging from 5 to 50. The evaluation metric is the top-*K* hitting ratio (HR@*K*). This metric is calculated as the number of shared sequences between the top-*K* predicted results and the ground truth, divided by *K*. On the synthetic dataset, LLMED-MAE, LLMED-combined and Tensor Sketch show nearly identical top performance across different values of *K*. However, on the real dataset, LLMED with triplet loss achieves the best results among all the metrics. MAE loss and combined loss variants of LLMED also significantly outperform other methods while the performance of Tensor Sketch shows a noticeable performance drop when transitioning from synthetic to the real data.

**Figure 6:**
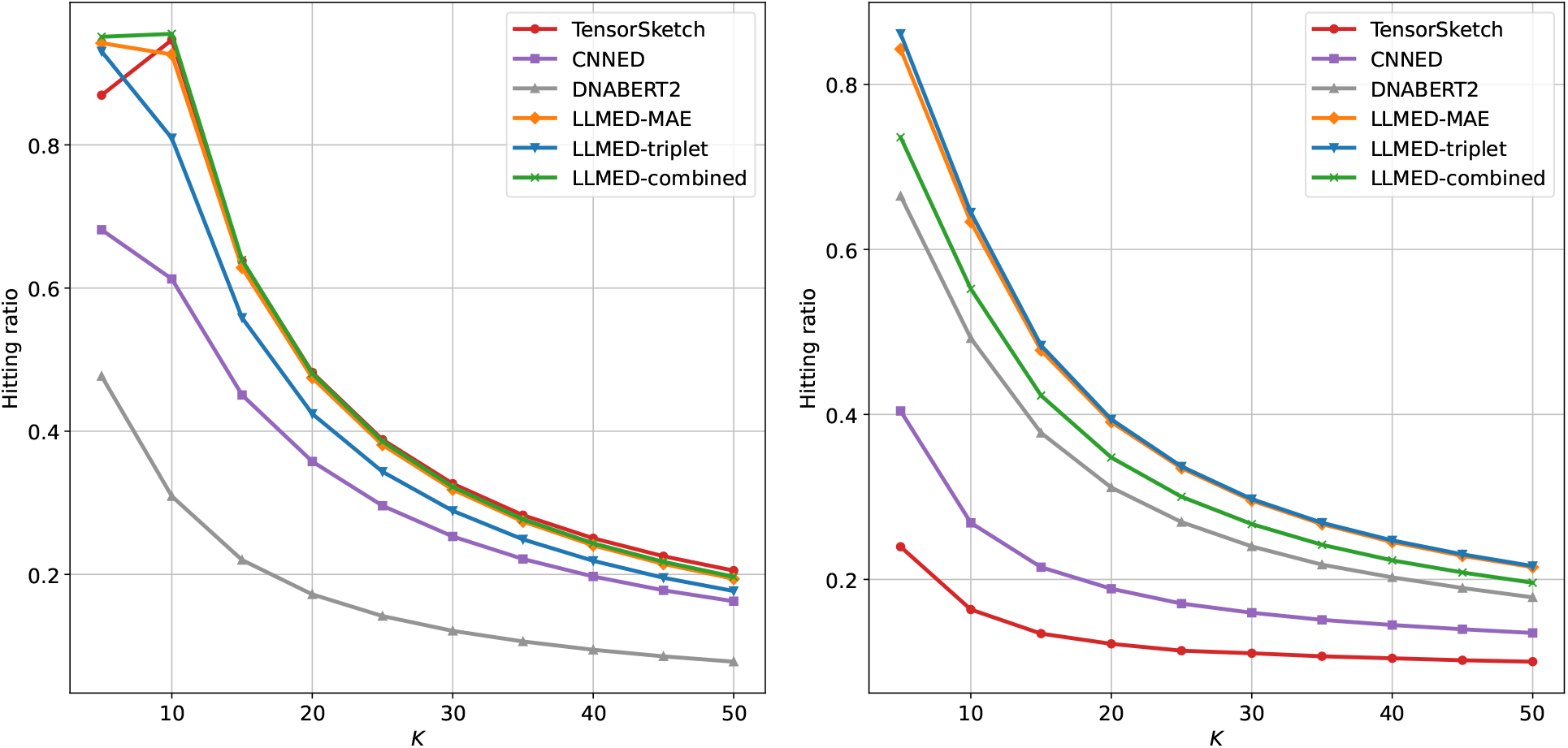
The hitting ratio for top-*K* on the synthetic dataset (left) and real dataset (right).

Fig. 7 and 8 illustrate the recall of top-1 and top-10 across two datasets, with varying numbers of predictions. The curves show that LLMED achieves the best performance across almost all measures. The only exception is the recall of top-10 on the synthetic dataset, where both LLMED and Tensor Sketch perform well. Moreover, LLMED with triplet loss demonstrates the best performance on Gen50ks, consistent with the results observed for the top-*K* hitting ratio. These findings suggest that the triplet loss effectively improves the model’s ability to generalize to more complex, real-world datasets with longer sequences.

**Figure 7:**
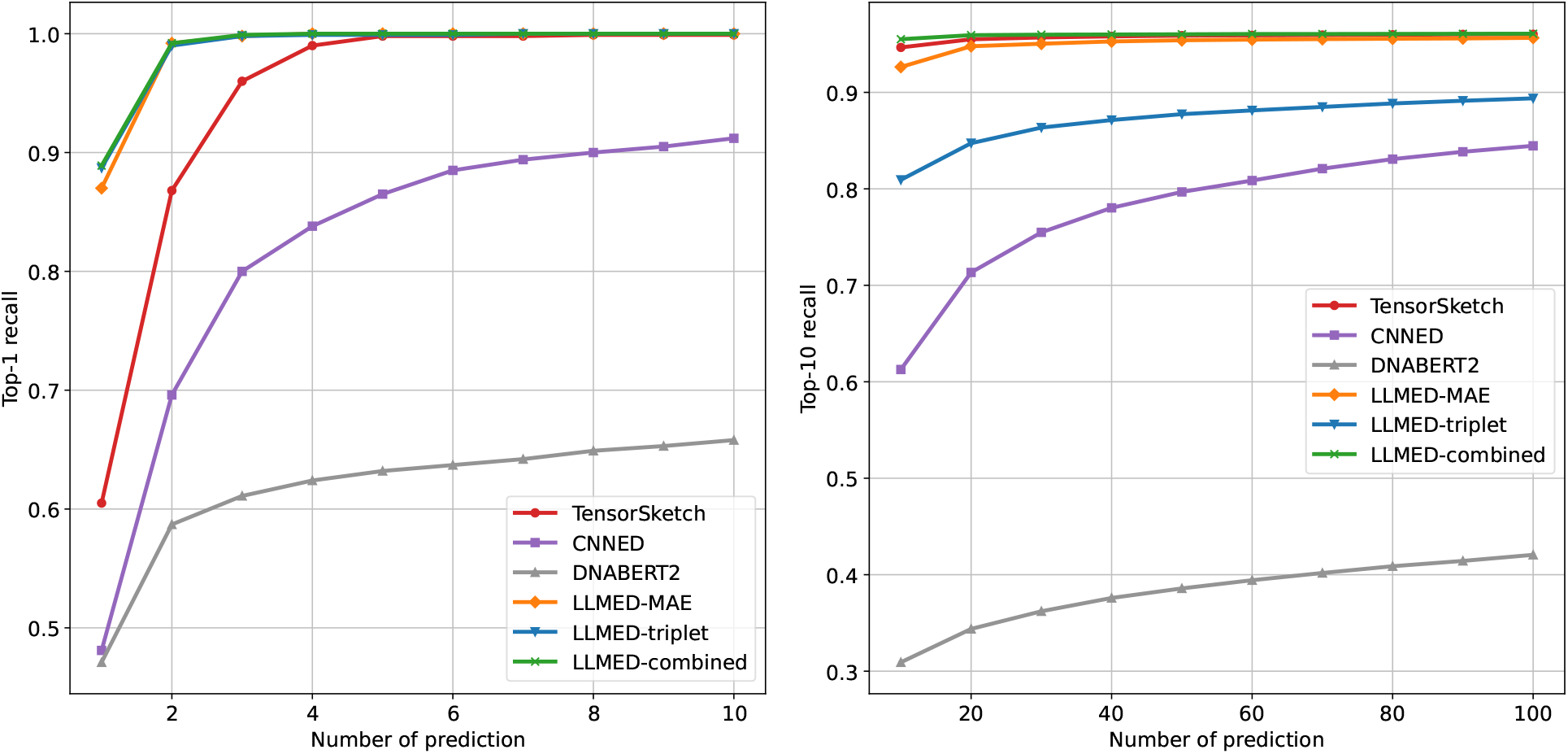
The recall of top-1 and top-10 on the synthetic dataset. The x-axis represents the number of prediction, and the y-axis represents the recall of top-*K*.

**Figure 8:**
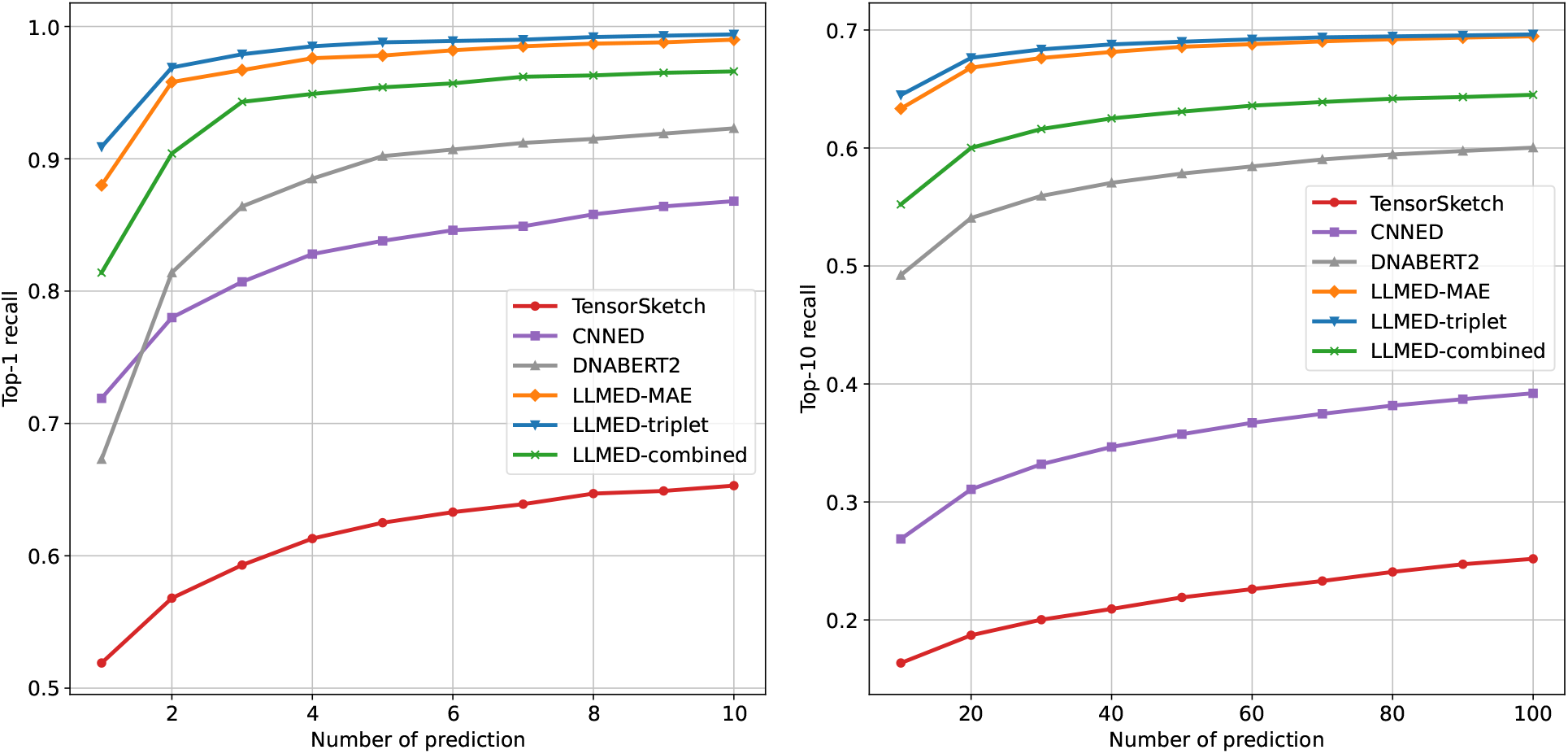
The recall of top-1 and top-10 on Gen50ks. The x-axis represents the number of prediction, and the y-axis represents the recall of top-*K*.

## 3 Conclusion

We propose LLMED, a novel genomic foundation model designed to produce sequence embeddings that approximate the edit distance. Our framework is versatile, enabling the training of LLMED using any existing genomic foundation models. It demonstrated the highest correlation with edit distance, affirming its superior embedding capabilities. In similar sequence search, our model also outperforms existing leading rule-based and machine-learning-based methods. As a result, LLMED shows great potential for applications involving edit distance-based sequence comparisons.

Edit distance embedding is a fundamental problem in computational biology. LLMED represents another successful deployment of genomic language models to tackle computational challenges in biology, shedding light on broader applications of genomic language models for solving algorithmic problems in biology.

LLMED can be enhanced by incorporating advanced techniques used in improving large language models, particularly those specifically designed to boost embedding quality in natural language processing. As an example, in [18], a model with additional latent attention layers has achieved state-of-the-art performance on benchmarked tasks. Adapting such methodologies to improve LLMED’s performance is a promising direction for our future work.

## 4 Methods

We first introduce the structure of LLMED, followed by the design of objective functions and training process.

### 4.1 Framework

The goal of our model is to embed genomic sequences, i.e., mapping sequences into a vector space. More specifically, given a sequence *S* over the alphabet Σ = {*A, C, G, T*}, our sequence embedding model *E* transforms *S* into a vector *E*(*S*) ∈ ℝ^*d*^, where *d* denotes the dimension of the embedding vector.

The framework of LLMED is shown in Fig. 9. Our framework uses a genomic LLM, and takes the output of the LLM to generate the sequence embedding. For a genomic LLM *F*, the input sequence *S* is first tokenized into a list of *n* tokens *x* = (*x*_1_, *x*_2_, …, *x*_*n*_). For each token, *F* generates a *d*-dimensional vector. Hence, *F* maps *S* into an *n* × *d* matrix, written as (*F* (*S*)_1_, *F* (*S*)_2_, …, *F* (*S*)_*n*_), where *F* (*S*)_*i*_ ∈ ℝ^*d*^, 1 ≤ *i* ≤ *n*. Our model *E* condenses this matrix into an embedding vector of dimension *d* by average-pooling. Formally,

**Figure 9:**
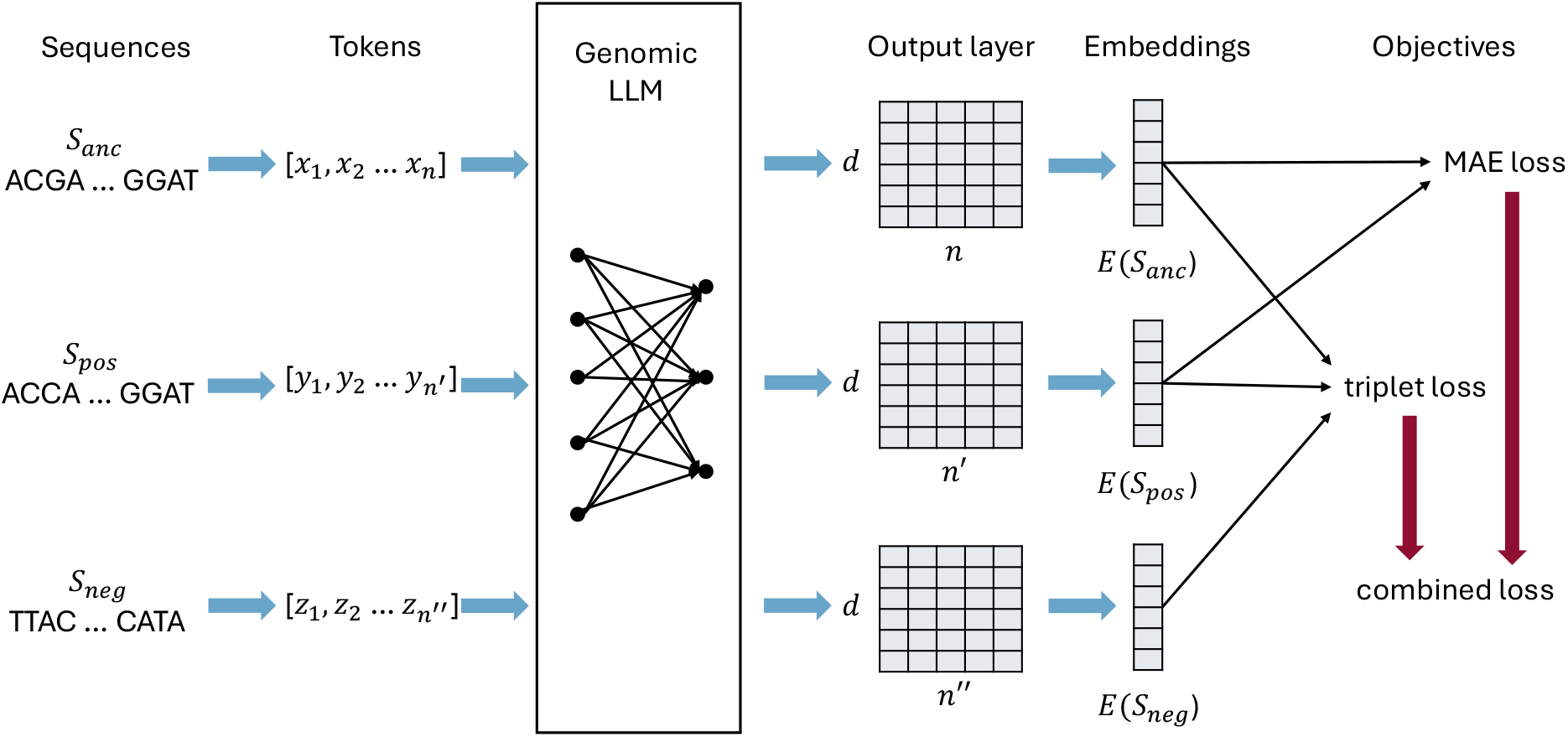
The architecture of LLMED.

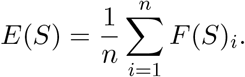

The utilized LLM can be any existing genomic foundation model. For this work, we choose to use DNABERT2, a transformer-based foundation model; we also use its tokenizer in our method.

### 4.2 Loss Functions

Our embedding model *E* is trained using contrastive learning; we explore three loss functions.

#### Mean Absolute Error Loss

This objective function approximates the edit distance between sequences by minimizing the mean absolute error (MAE) between the cosine similarity of the embeddings and the edit similarity of the input sequences (as defined below). Doing so ensures that the embedding space accurately reflects sequence similarity. Formally, let (*S*_1_, *S*_2_) be a pair of sequences; let *s*(*S*_1_, *S*_2_) be their edit similarity, defined as

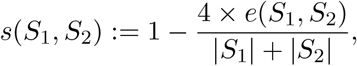

where *e*(*S*_1_, *S*_2_) is the edit distance between *S*_1_ and *S*_2_; |*S*_1_| and |*S*_2_| represent the lengths of *S*_1_ and *S*_2_, respectively. Note that *s*(*S*_1_, *S*_2_) is scaled to a desired range of [−1, 1].

The loss over this pair of sequences is defined as:

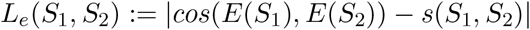

where 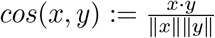 is the cosine similarity between *x* and *y*.

Let S = {(*S*_1_, *S*_2_)} be the training set consisting of multiple sequence pairs. The MAE loss is the average loss over all individual pairs:

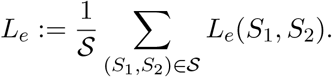

#### Triplet Loss

This objective utilizes triplets of sequences: an anchor sequence *S*_*a*_, a positive sequence *S*_*p*_, and a negative sequence *S*_*n*_. We require that every training sample (*S*_*a*_, *S*_*p*_, *S*_*n*_) satisfies the property that the edit distance between *S*_*a*_ and *S*_*p*_ is strictly less than that between *S*_*a*_ and *S*_*n*_. The triplet loss over one training sample is defined as:

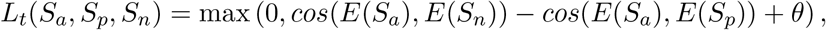

where *θ* is a margin hyperparameter. Similarly, the total triplet loss *L*_*t*_ is the averaged triplet loss over all training samples.

#### Combined Loss

Let (*S*_*a*_, *S*_*p*_, *S*_*n*_) be a training sample. The combined loss takes into account both the triplet loss and the MAE loss between the anchor sequence *S*_*a*_ and the positive sequence *S*_*p*_. Formally,

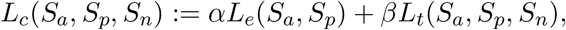

where *α* and *β* are two weight hyperparameters. The total combined loss *L*_*c*_ is the average over all training samples.

### 4.3 Training Data and Procedure

#### Mean Absolute Error Loss

For MAE loss, we simulate one million pairs of sequences. Each pair consists of a randomly generated sequence of length *L* over the alphabet Σ = {*A, C, G, T*}, and a second sequence obtained by applying random mutations to the first one. The mutation rate is sampled uniformly from 3% to 30%, with mutations equally distributed among substitutions, insertions, and deletions. The dataset is divided into two parts, with 95% as training dataset and 5% as validation dataset. Training starts from the pretrained DNABERT2 model (DNABERT-2-117M), and runs for 10 epochs with a learning rate of 10^−5^ and a batch size of 16. Checkpoints are saved every 5000 training steps, and the checkpoint with the lowest validation loss is selected as the final model state.

#### Triplet Loss

For training with the triplet loss, each data sample consists of three sequences: an anchor sequence, a positive sequence, and a negative sequence. Similar to the MAE dataset, the anchor is a randomly generated sequence of length *L*. The positive sequence is generated by randomly applying mutations on the anchor sequence with 10% mutation rate. And the negative sequence is generated similarly but with a higher mutation rate of 40%. The choice of these mutation rates are based on the observation that two random sequences typically have a similarity of around 50%. By choosing a slightly lower mutation rate for the negative sequence, we aim to enable the model to distinguish biologically related or similar sequences from dissimilar (yet not totally random) sequences more effectively. The dataset is split into a 90% training set and a 10% validation set. The margin hyperparameter *θ* is set to 0.3, and other training settings are identical to those for the MAE loss.

#### Combined Loss

The dataset used for combined loss is the same as for the triplet loss. The weighting hyperparameters are set to *α* = 0.5 and *β* = 1. The other training settings remain consistent with the triplet loss training. The curves of training and validation losses during training are shown in Fig. 10.

**Figure 10:**
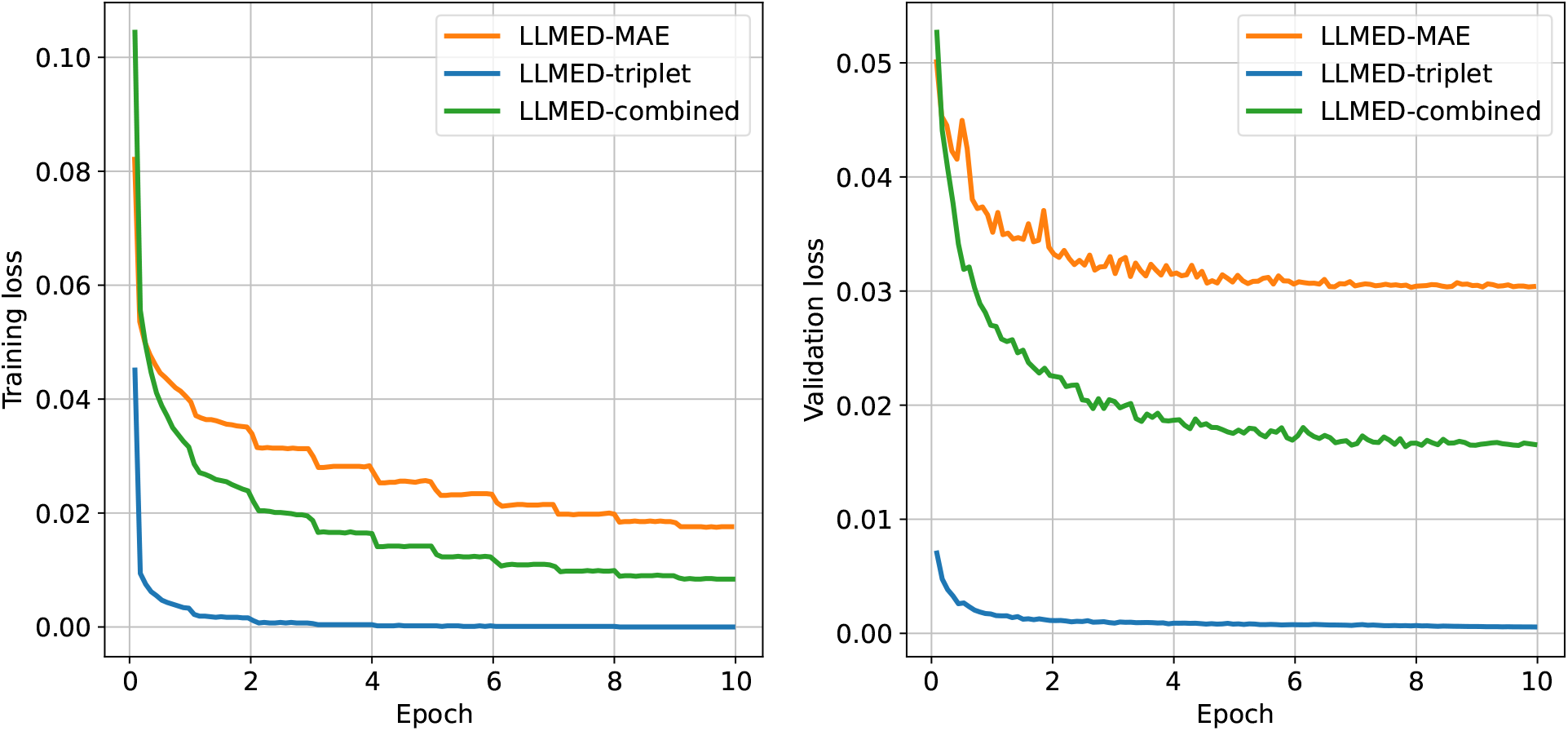
Left: The training loss of the three loss functions during training. Right: The validation loss of the three loss functions during training.

## Availability of data and materials

The code for LLMED is available on GitHub (https://github.com/Shao-Group/llmembedding). Data is available at: https://doi.org/10.5281/zenodo.17161249.

## Authors’ contributions

X.L. designed and implemented the method, conducted the majority of the experiments, and performed the computational analyses. K.C. implemented the similarity search pipeline and contributed to the analysis of the similarity search results. Y.Z. implemented some of the baseline methods and contributed to the presentation of the corresponding figures. M.S. supervised the study. All authors contributed to the writing of the manuscript and approved the final version.

## Acknowledgments

Not applicable

## References

[1] A. Andoni, M. Deza, A. Gupta, P. Indyk, and S. Raskhodnikova. Lower bounds for embedding edit distance into normed spaces. In Proceedings of the Fourteenth Annual ACM-SIAM Symposium on Discrete Algorithms, SODA ’03, page 523–526, USA, 2003. Society for Industrial and Applied Mathematics.

[2] Z. Bar-Yossef, T.S. Jayram, R. Krauthgamer, and R. Kumar. Approximating edit distance efficiently. In 45th Annual IEEE Symposium on Foundations of Computer Science, pages 550–559, 2004.

[3] Tuğkan Batu, Funda Ergun, and Cenk Sahinalp. Oblivious string embeddings and edit distance approximations. In Proceedings of the Seventeenth Annual ACM-SIAM Symposium on Discrete Algorithm, SODA ’06, page 792–801, USA, 2006. Society for Industrial and Applied Mathematics.

[4] A.Z. Broder. On the resemblance and containment of documents. In Proceedings. Compression and Complexity of SEQUENCES 1997 (Cat. No.97TB100171), pages 21–29, 1997.

[5] Diptarka Chakraborty, Elazar Goldenberg, and Michal Koucký. Streaming algorithms for embedding and computing edit distance in the low distance regime. In Proceedings of the Forty-Eighth Annual ACM Symposium on Theory of Computing, STOC ’16, page 712–725, New York, NY, USA, 2016. Association for Computing Machinery.

[6] Zhihao Chang, Linzhu Yu, Yanchao Xu, and Wentao Hu. Neural embeddings for knn search in biological sequence. Proceedings of the AAAI Conference on Artificial Intelligence, 38(1):38–45, Mar. 2024.

[7] Gabriele Corso, Zhitao Ying, Michal Pándy, Petar Veličković, Jure Leskovec, and Pietro Lio. Neural distance embeddings for biological sequences. In M. Ranzato, A. Beygelzimer, Y. Dauphin, P.S. Liang, and J. Wortman Vaughan, editors, Advances in Neural Information Processing Systems, volume 34, pages 18539–18551. Curran Associates, Inc., 2021.

[8] Xinyan Dai, Xiao Yan, Kaiwen Zhou, Yuxuan Wang, Han Yang, and James Cheng. Convolutional embedding for edit distance. In Proceedings of the 43rd International ACM SIGIR Conference on Research and Development in Information Retrieval, SIGIR ’20, page 599–608, New York, NY, USA, 2020. Association for Computing Machinery.

[9] Hugo Dalla-Torre, Liam Gonzalez, Javier Mendoza-Revilla, Nicolas Lopez Carranza, Adam Henryk Grzywaczewski, Francesco Oteri, Christian Dallago, Evan Trop, Bernardo P. de Almeida, Hassan Sirelkhatim, Guillaume Richard, Marcin Skwark, Karim Beguir, Marie Lopez, and Thomas Pierrot. Nucleotide transformer: building and evaluating robust foundation models for human genomics. Nature Methods, Nov 2024.

[10] Jacob Devlin, Ming-Wei Chang, Kenton Lee, and Kristina Toutanova. BERT: Pre-training of deep bidirectional transformers for language understanding. In Jill Burstein, Christy Doran, and Thamar Solorio, editors, Proceedings of the 2019 Conference of the North American Chapter of the Association for Computational Linguistics: Human Language Technologies, Volume 1 (Long and Short Papers), pages 4171–4186, Minneapolis, Minnesota, June 2019. Association for Computational Linguistics.

[11] Robert Edgar. Syncmers are more sensitive than minimizers for selecting conserved k-mers in biological sequences. PeerJ, 9:e10805, 2021.

[12] Ragnar Groot Koerkamp. A*pa2: up to 20 times faster exact global alignment. bioRxiv, 2024.

[13] Albert Gu and Tri Dao. Mamba: Linear-time sequence modeling with selective state spaces. arXiv preprint arXiv:2312.00752, 2023.

[14] Chirag Jain, Alexander Dilthey, Sergey Koren, Srinivas Aluru, and Adam M. Phillippy. A fast approximate algorithm for mapping long reads to large reference databases. Journal of Computational Biology, 25(7): 766–779, 2018. PMID: 29708767.

[15] Yanrong Ji, Zhihan Zhou, Han Liu, and Ramana V Davuluri. Dnabert: pre-trained bidirectional encoder representations from transformers model for dna-language in genome. Bioinformatics, 37(15):2112–2120, 02 2021.

[16] Amir Joudaki, Alexandru Meterez, Harun Mustafa, Ragnar Groot Koerkamp, André Kahles, and Gunnar Rätsch. Aligning distant sequences to graphs using long seed sketches. Genome Research, 33(7): 1208–1217, 2023.

[17] Amir Joudaki, Gunnar Rätsch, and André Kahles. Fast alignment-free similarity estimation by tensor sketching. bioRxiv, 2021.

[18] Chankyu Lee, Rajarshi Roy, Mengyao Xu, Jonathan Raiman, Mohammad Shoeybi, Bryan Catanzaro, and Wei Ping. Nv-embed: Improved techniques for training llms as generalist embedding models, 2025.

[19] Matthew Macaulay, Aaron Darling, and Mathieu Fourment. Fidelity of hyperbolic space for bayesian phylogenetic inference. PLOS Computational Biology, 19(4):1–20, 04 2023.

[20] Santiago Marco-Sola, Juan Carlos Moure, Miquel Moreto, and Antonio Espinosa. Fast gap-affine pairwise alignment using the wavefront algorithm. Bioinformatics, 37(4):456–463, 09 2020.

[21] Eric Nguyen, Michael Poli, Matthew G. Durrant, Brian Kang, Dhruva Katrekar, David B. Li, Liam J. Bartie, Armin W. Thomas, Samuel H. King, Garyk Brixi, Jeremy Sullivan, Madelena Y. Ng, Ashley Lewis, Aaron Lou, Stefano Ermon, Stephen A. Baccus, Tina Hernandez-Boussard, Christopher Ré, Patrick D. Hsu, and Brian L. Hie. Sequence modeling and design from molecular to genome scale with evo. Science, 386(6723):eado9336, 2024.

[22] Eric Nguyen, Michael Poli, Marjan Faizi, Armin W. Thomas, Callum Birch Sykes, Michael Wornow, Aman Patel, Clayton Rabideau, Stefano Massaroli, Yoshua Bengio, Stefano Ermon, Stephen A. Baccus, and Christopher Ré. Hyenadna: long-range genomic sequence modeling at single nucleotide resolution. In Proceedings of the 37th International Conference on Neural Information Processing Systems, NIPS ’23, Red Hook, NY, USA, 2024. Curran Associates Inc.

[23] Rafail Ostrovsky and Yuval Rabani. Low distortion embeddings for edit distance. J. ACM, 54(5):23–es, October 2007.

[24] Mahmudur Rahman Hera, Paul Medvedev, David Koslicki, and Antonio Blanca. Estimation of Substitution and Indel Rates via k-mer Statistics. In Broňa Brejová and Rob Patro, editors, 25th International Conference on Algorithms for Bioinformatics (WABI 2025), volume 344 of Leibniz International Proceedings in Informatics (LIPIcs), pages 16:1–16:15, Dagstuhl, Germany, 2025. Schloss Dagstuhl – Leibniz-Zentrum für Informatik.

[25] Michael Roberts, Wayne Hayes, Brian R. Hunt, Stephen M. Mount, and James A. Yorke. Reducing storage requirements for biological sequence comparison. Bioinformatics, 20(18):3363–3369, 07 2004.

[26] Kristoffer Sahlin. Effective sequence similarity detection with strobemers. Genome research, 31(11): 2080–2094, 2021.

[27] Kristoffer Sahlin. Strobealign: flexible seed size enables ultra-fast and accurate read alignment. Genome Biology, 23(1): 260, 2022.

[28] Saul Schleimer, Daniel S. Wilkerson, and Alex Aiken. Winnowing: local algorithms for document fingerprinting. In Proceedings of the 2003 ACM SIGMOD International Conference on Management of Data, SIGMOD ’03, page 76–85, New York, NY, USA, 2003. Association for Computing Machinery.

[29] Jim Shaw and Yun William Yu. Fast and robust metagenomic sequence comparison through sparse chaining with skani. Nature Methods, 20(11): 1661–1665, 2023.

[30] Gregory E. Sims, Se-Ran Jun, Guohong A. Wu, and Sung-Hou Kim. Alignment-free genome comparison with feature frequency profiles (ffp) and optimal resolutions. Proceedings of the National Academy of Sciences, 106(8): 2677–2682, 2009.

[31] Yan Song, Haixu Tang, Haoyu Zhang, and Qin Zhang. Overlap detection on long, error-prone sequencing reads via smooth q-gram. Bioinformatics, 36(19):4838–4845, 04 2020.

[32] Ashish Vaswani, Noam Shazeer, Niki Parmar, Jakob Uszkoreit, Llion Jones, Aidan N Gomez, Lukasz Kaiser, and Illia Polosukhin. Attention is all you need. In I. Guyon, U. Von Luxburg, S. Bengio, H. Wallach, R. Fergus, S. Vishwanathan, and R. Garnett, editors, Advances in Neural Information Processing Systems, volume 30. Curran Associates, Inc., 2017.

[33] Haoyu Zhang and Qin Zhang. Embedjoin: Efficient edit similarity joins via embeddings. In Proceedings of the 23rd ACM SIGKDD International Conference on Knowledge Discovery and Data Mining, KDD ’17, page 585–594, New York, NY, USA, 2017. Association for Computing Machinery.

[34] Xiyuan Zhang, Yang Yuan, and Piotr Indyk. Neural embeddings for nearest neighbor search under edit distance, 2020.

[35] Zhihan Zhou, Yanrong Ji, Weijian Li, Pratik Dutta, Ramana V Davuluri, and Han Liu. DNABERT-2: Efficient foundation model and benchmark for multi-species genomes. In The Twelfth International Conference on Learning Representations, 2024.

[36] Martin Šošić and Mile Šikić. Edlib: a c/c++ library for fast, exact sequence alignment using edit distance. Bioinformatics, 33(9):1394–1395, 01 2017.

